# A simple action reduces high fat diet intake and obesity in mice

**DOI:** 10.1101/2024.10.01.615599

**Authors:** M. R. Barrett, Y. Pan, Chantelle Murrell, Eva O. Karolczak, Justin Wang, Lisa Fang, Jeremy M. Thompson, Yu-Hsuan Chang, Eric Casey, J. Czarny, Wang Lok So, Alex Reichenbach, Romana Stark, Hamid Taghipourbibalan, Suzanne R. Penna, Katherine B. McCullough, Sara Westbrook, Bridget Matikainen-Ankney, Victor A Cazares, Kristen Delevich, Wambura Fobbs, Susan Maloney, Ames Sutton Hickey, James E. McCutcheon, Zane Andrews, Meaghan C. Creed, Michael J. Krashes, Alexxai V. Kravitz

**Author notes:** These authors contributed equally to this work (M. R. Barrett, Y. Pan).

## Abstract

Diets that are high in fat cause over-eating and weight gain in multiple species of animals, suggesting that high dietary fat is sufficient to cause obesity. However, high-fat diets are typically provided freely to animals in obesity experiments, so it remains unclear if high-fat diets would still cause obesity if they required more effort to obtain. We hypothesized that unrestricted and easy access is necessary for high-fat diet induced over-eating, and the corollary that requiring mice to perform small amounts of work to obtain high-fat diet would reduce high-fat diet intake and associated weight gain. To test this hypothesis, we developed a novel home-cage based feeding device that either provided high-fat diet freely, or after mice poked their noses into a port one time – a simple action that is easy for them to do. We tested the effect of this intervention for six weeks, with mice receiving all daily calories from high-fat diet, modifying only how they accessed it. Requiring mice to nose-poke to access high-fat diet reduced intake and nearly completely prevented the development of obesity. In follow up experiments, we observed a similar phenomenon in mice responding for low-fat grain-based pellets that do not induce obesity, suggesting a general mechanism whereby animals engage with and consume more food when it is freely available vs. when it requires a simple action to obtain. We conclude that unrestricted access to food promotes overeating, and that a simple action such as a nose-poke can reduce over-eating and weight gain in mice. This may have implications for why over-eating and obesity are common in modern food environments, which are often characterized by easy access to low-cost unhealthy foods.

## Introduction

The United States Department of Agriculture (USDA) tracks historical trends in food availability (1), revealing that total available calories per capita in the US rose by about 23% between 1970 and 2014, accounted for primarily by changes in available grains and fats (2,3). While the USDA data does not quantify food consumption over this period, an orthogonal dataset utilizing dietary recall, the National Health and Nutrition Examination Survey (NHANES), reported a 12-15% increase in calorie consumption between 1970 to 2010 (4). While increases in food consumption are likely driving the obesity epidemic (5,6), questions still remain: Why are Americans eating more than they were eating in 1970? Has something changed in our food supply or environment that is causing us to overeat? Possible explanations include ubiquitous access to low-cost unhealthy foods (7,8), changes in nutrient contents and processing (5), interactions between genetic predispositions and the modern food supply (9), chemicals in the environment including those that disrupt endocrine function (10,11), disruptions in sleep and circadian rhythms (12), and other causes (9,13). As it is challenging to isolate specific factors in humans, laboratory animals have often been used to test how specific factors alter body weight.

Increasing dietary fat content induces over-eating and weight gain in multiple species, including monkeys, dogs, pigs, hamsters, squirrels, rats, and mice (14,15), and has been used for more than 75 years to model obesity in animals (16). The face validity of the “high-fat diet” model of obesity relies on observational studies showing that higher dietary fat consumption is also associated with higher obesity prevalence in humans (17,18), and interventional studies demonstrating that low-fat diets produce modest, but significant, weight loss in humans (19,20). However, in animal models the high-fat diets are typically placed in a food hopper in the animal’s cage, providing unrestricted easy access. While this may replicate the extremes of how ubiquitous snack foods have become in modern life, it fails to test whether such unrestricted access is a necessary component for inducing high-fat diet induced obesity. We hypothesized that unrestricted and easy access is necessary for high-fat diet induced over-eating, and the corollary that requiring mice to perform small amounts of work to obtain high-fat diet would reduce high-fat diet intake and associated weight gain.

We designed a novel home-cage feeding device, the Tumble Feeder, to test these hypotheses. The Tumble Feeder contains a moveable food hopper that can provide access to high-fat diet freely by leaving the hopper open, or in a controlled fashion that requires the mouse to touch a trigger with its nose to open the hopper (termed: a nosepoke). We found that requiring mice to perform a single nose-poke greatly reduced daily intake of high-fat diet, and almost completely blocked associated weight gain. This was surprising as the nose-poke action does not require strong physical effort to perform, and mice could earn as much high-fat diet as they wanted each day by doing more nose-pokes. We performed analogous follow-up experiments to test if our results were specific to high-fat diet or reflected a more general behavioral control over food intake. Here, we observed that the nose-poke requirement also reduced how many low-fat grain-based pellets were taken, which was attributed most strongly to a reduction in the number of feeding bouts. This suggests that the constant presence of food induced mice to initiate additional bouts of feeding each day, which can explain their over-eating and weight gain. Together, our experiments support our hypothesis that a small action – one nose-poke – can reduce food intake and diet-induced weight gain, and that easy access to unhealthy foods may be a critical contributor to the human obesity epidemic.

## Results

### An operant device to control access to high-fat diet

We developed a novel operant device that allowed us to control access to high-fat diet over multiple weeks in the home-cage (the TumbleFeeder, design files available at https://github.com/KravitzLabDevices/CastleFeeder/, Fig 1A-D). The Tumble Feeder has two touch-sensitive nose-poke triggers for detecting mouse “nose-pokes”, touch sensitive bars to detect food hopper interactions, a microcontroller, screen, real-time clock (RTC), and a micro secure-digital (microSD) card slot for controlling tasks and displaying and logging data (Fig 1B, C). We first programmed the Tumble Feeder so the hopper was either open for free access (ie: Free mode) or opened for 60s every time the mouse touched a nose-poke (ie: FR1 mode). In both Free and FR1 modes, mice were allowed to access the hopper an unlimited number of times per day, with no imposed delays between nose-pokes in the FR1 mode.

**Figure 1:**
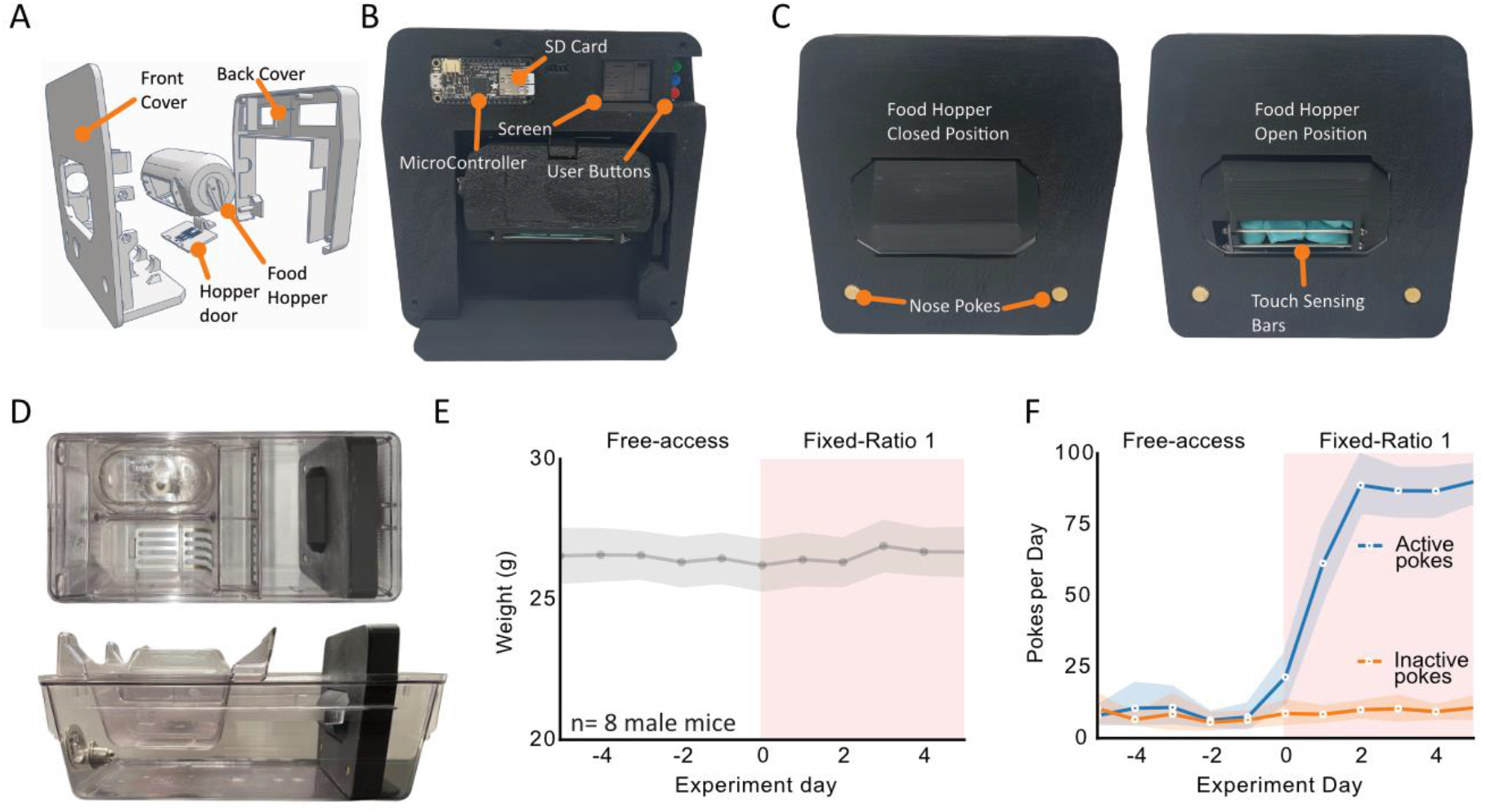
Design of the Tumble Feeder. **A**. 3D design of the Tumble Feeder. **B**. Back view of the assembled Tumble Feeder with relevant part labels. **C**. Front view of the Tumble Feeder in both open- and closed positions. **D**. Photographs of Tumble Feeder in a mouse home-cage. **E**. Average daily weight of mice during Free vs. FR1 tasks for laboratory chow (n=8). **F**. Number of active and inactive pokes per day in the Free and FR1 modes (n=8 mice, 5 days in each phase).

To validate the performance of the Tumble Feeder, 8 male C57Bl6 mice were given access to the Free task for 5 days, with laboratory chow in the feeder. The Tumble Feeders were then switched to FR1 mode for an additional 5 days, with the same diet. To control for the movement of the hopper possibly startling the mice in the FR1 phase, we programmed the hopper to open and close every 15 minutes throughout the Free phases, resulting in 96 actuations per day, slightly higher than the number of openings in the FR1 phase. Mice rapidly learned to operate the Tumble Feeder on FR1, as evidenced by an increasing number of pokes per day (Fig 1F). Critically, the nose-poke is an easy action to complete, and each nose-poke opened the hopper for 60s, which was long enough for mice to easily obtain their daily caloric needs each day. As such, mice maintained their body weights during both tasks, confirming that mice readily learned how to operate the Tumble Feeder in the FR1 mode, and could obtain their necessary daily caloric requirements in both the Free and FR1 modes (Fig 1E).

### A single nose-poke can reduce high-fat diet intake and associated weight gain

To test if requiring mice to nose-poke to access high-fat diet would reduce intake and weight gain, 12 C57Bl6 male mice were first given access to the Tumble Feeder for 5 days with laboratory chow in the hopper. This allowed the mice to acclimate to the Tumble Feeder and allowed us to quantify daily intake of chow before starting the high-fat diet experiment. Next, the Tumble Feeder was filled with high-fat diet (60% of calories from fat, Research Diets #D12492) and programmed to alternate in 5-day phases of Free and FR1 modes, with each nose-poke providing 60s of access to high-fat diet in the FR1 phases (Fig 2A, B, Movie S1).

**Figure 2:**
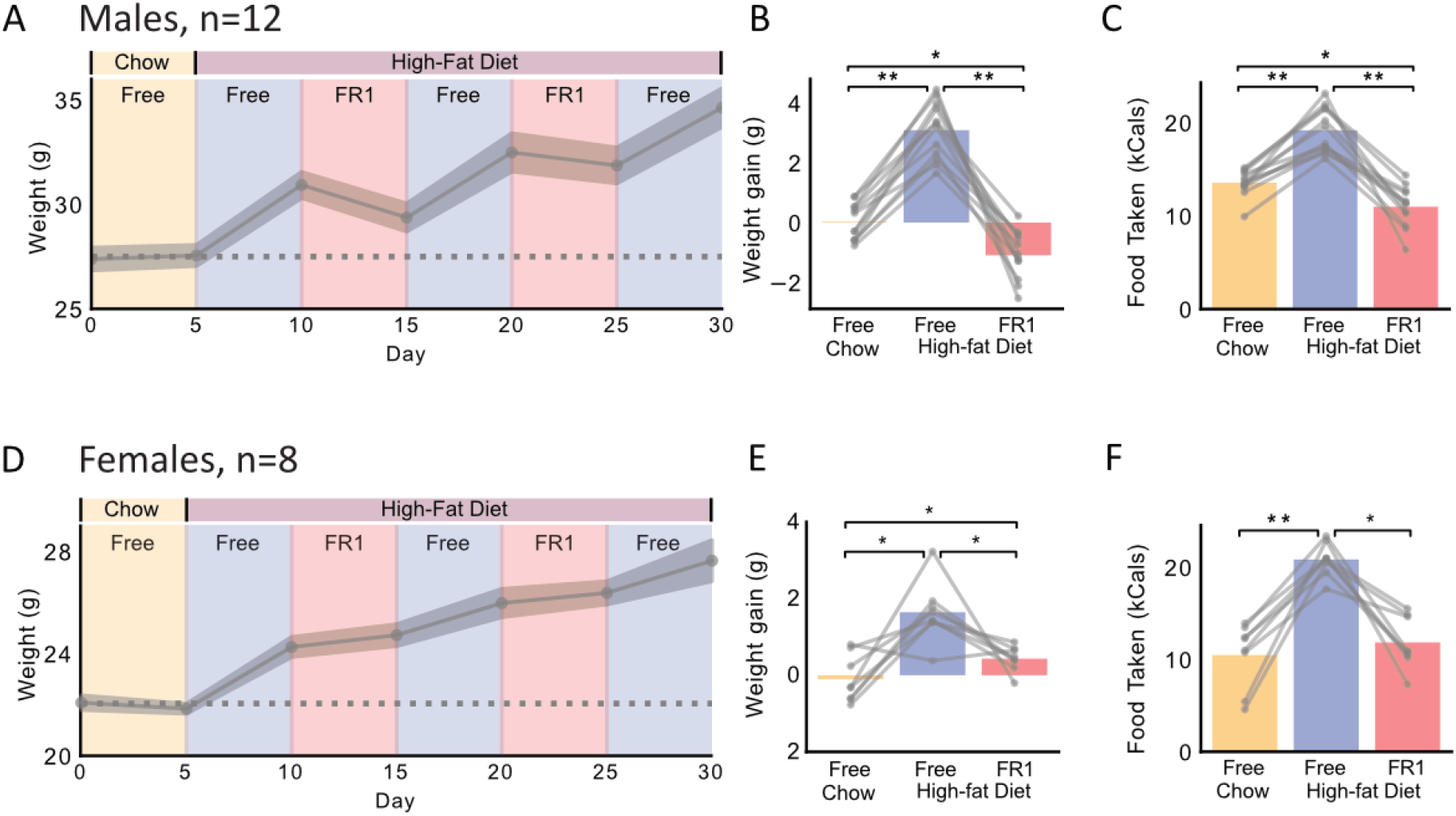
Mice take less high-fat diet and gain less weight on FR1 vs. Free access. **A**. Average weight of mice across Free and FR1 phases for chow and high-fat diet (n=12 male mice). **B**. Weight gain in each phase. **C**. Food taken in each phase. **D-F**. same plot format as **A-C**, but for 8 female mice. ** denotes p<0.001 * denotes p<0.05.

Consistent with our primary hypothesis, mice only gained weight in the Free high-fat diet phases and lost a small amount of weight in the FR1 high-fat diet phases (Free: +3.1g / 5 days, FR1: -1.1g / 5 days, significant effect of Task: F(2, 22) = 85.9, p < 0.001, post-hoc paired t-test between Free and FR1, p < 0.001, Fig 2A, B). The Tumble Feeders were also weighed throughout the experiment to quantify how much high-fat diet was removed in each phase.

Consistent with the difference in weight gain, mice removed significantly more high-fat diet in the Free high-fat diet phase than the FR1 high-fat diet phase (Free: 19kcal/day vs. FR1: 11kcal/day, paired t-test p<0.001, Fig 2C). We repeated this experiment with 8 female mice, who also gained significantly more weight in Free high-fat diet than FR1 high-fat diet phases (Free: +1.7g / 5 days, FR1: +0.4g / 5 days, significant effect of Task: F(2, 14) = 12.63, p < 0.001, post-hoc paired t-test between Free and FR1, p =0.01, Fig 2D, E), and removed significantly more high-fat diet in the Free phase vs. the FR1 phase (Free: 21kcal/day vs. FR1: 12kcal/day, paired t-test p=0.001, Fig 2F).

### FR1 access to high-fat diet prevents obesity

We next tested if requiring a nose-poke to obtain high-fat diet would prevent the development of obesity over 6 weeks of high-fat diet exposure. Male mice were used in this experiment as female mice are more often resistant to high-fat diet induced weight gain (21). Seven new male C57Bl6 mice were given high-fat diet on the FR1 task for 6 weeks. Despite eating 100% of their daily calories from high-fat diet, these mice exhibited minimal weight gain over the 6 weeks, which did not differ significantly from the expected growth curve of age-matched male C57Bl6 mice (data from Jax, dashed line on Fig 3A, one-sample t-test p =0.138). Importantly, there was no restriction on how much high-fat diet mice could earn each day, and the average number of nose-pokes per day was ∼50, meaning mice opened the hopper for an average of ∼50 minutes per day to obtain their daily calories. As such, there was also ample time (>23 hours per day) for the mice to nose-poke more to obtain more high-fat diet if they had wanted to. At the conclusion of this experiment, the Tumble Feeders were switched to the Free mode for 6 additional weeks, resulting in a rapid increase in body weight (Fig 3A, B), confirming that the FR1 task was suppressing high-fat diet-induced weight gain in these mice (weight gain in FR1: 7.9 grams, in Free: 19.4 grams, paired t-test p<0.001). This supports our hypothesis that a necessary component of the high-fat diet-induced obesity paradigm is unrestricted easy access to high-fat diet, and requiring mice to perform a single nose-poke to access high-fat diet is sufficient to prevent weight gain.

**Figure 3:**
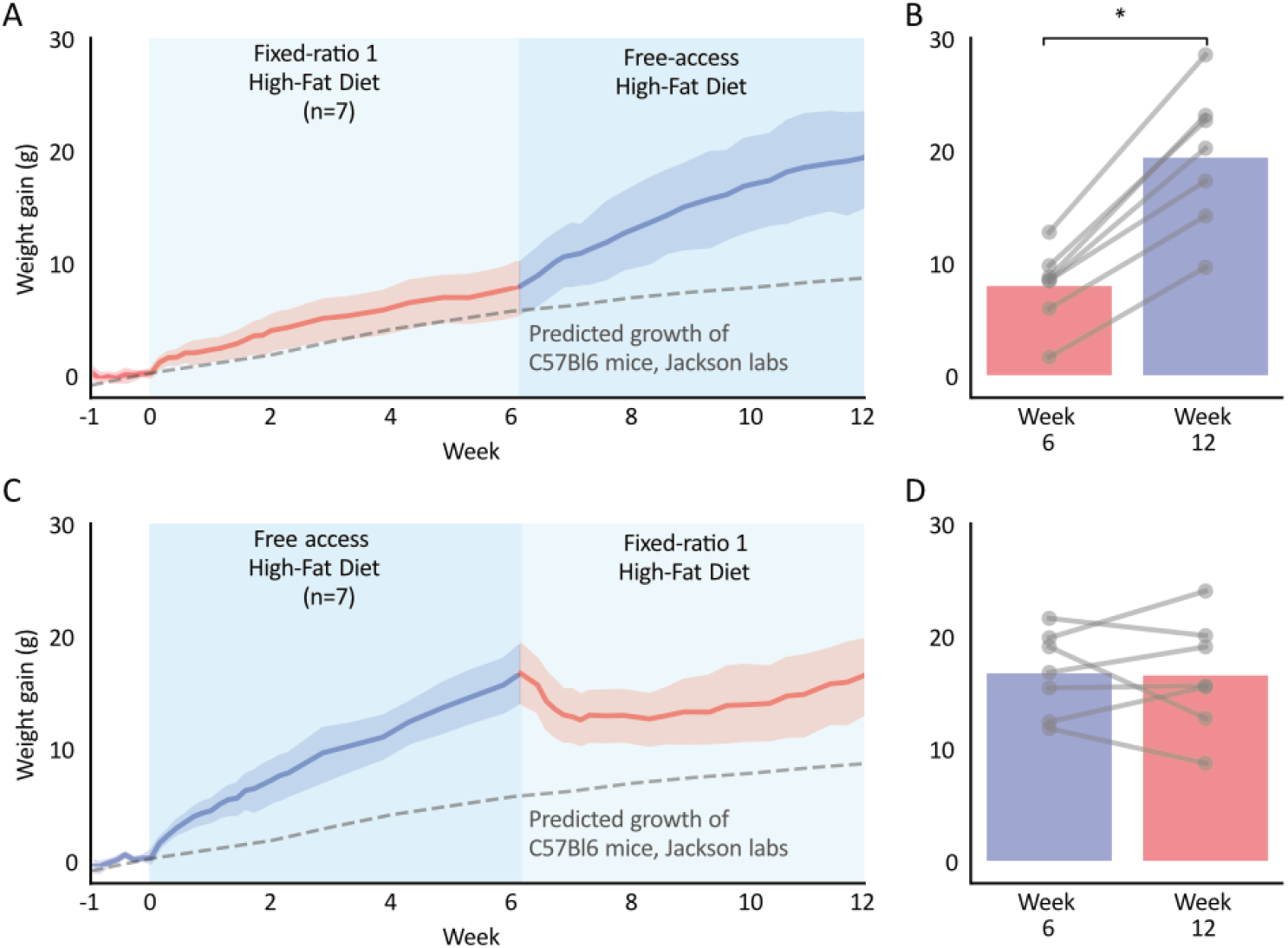
FR1 access to high-fat diet prevents, but does not reverse, obesity. **A**. Weight of mice on FR1 for high fat diet for 6 weeks, followed by Free access for six weeks (n=7 male mice). Dashed line is predicted growth curve of age-matched C57Bl6 mice from Jackson labs. **B**. Average weight at the end of the FR1 (week 6) vs. Free periods (week 12). **C**. Average weight of mice on Free access for high fat diet for 6 weeks, followed by FR1 access for six weeks (n=7 male mice). **D**. Average weight at the end of the Free (week 6) vs. FR1 access period (week 12). ** denotes p<0.001 * denotes p<0.05.

### Requiring mice to nose-poke for high-fat diet does not cause weight loss in obese mice

We next tested if requiring a nose-poke to access high-fat diet would result in weight loss in obese mice. Here, 7 new male C57Bl6 mice ate high-fat diet from the Tumble Feeder in Free mode for 6 weeks, resulting in weight gain (Fig 3C). The Tumble Feeders were then switched to FR1 for 6 additional weeks. While the mice lost a small amount of weight in the first week of FR1, they regained this weight over the next 6 weeks, resulting in a similar average weight at the start and end of the FR1 phase (weight gain in Free: 16.7 grams, FR1: 16.5 grams paired t-test, p = 0.90, Fig 3D). We conclude that the FR1 nose-poke requirement does not drive weight loss in obese mice. This highlights how the mechanisms that control the development of obesity can not necessarily be reversed to drive weight loss.

### Mice take fewer low-fat grain pellets when they have to nose-poke to access them

We hypothesized that the protective effect of the nose-poke requirement on high-fat diet induced obesity may not be specific to high-fat diets, but instead reflect a more general property of food intake, where animals over-eat when food is freely available vs. when it requires a small effort to obtain. To test this idea, we used Feeding Experimentation Device version 3 (FED3), a smart pellet dispensing device that operates in the home-cage (22). An advantage of using the FED3 over Tumble Feeder for these experiments was that we could obtain more detailed information about feeding patterns of mice with FED3. FED3 was programmed to operate in either a free-feeding mode (Free) or a fixed-ratio 1 (FR1) mode, to mimic the modes used in our high-fat diet experiments with the Tumble Feeder. In the Free mode, 20mg grain-based food pellets (Bio-serv F0163) were freely available in the pellet well and were replaced automatically whenever a mouse removed one. In the FR1 mode, the mouse had to break a photo-beam sensor with its nose (termed: a nose-poke) to cause FED3 to dispense each 20mg food pellet. There was no limit on how many pellets mice could obtain in either task and in both tasks FED3 was their only source of food.

We crowd sourced this experiment across multiple labs that use FED3, collecting Free and FR1 pellet data from 105 mice (72M/33F) across 10 labs in three countries (Fig 4A). All mice included in the analysis completed at least one day of Free followed by at least one day of FR1, and the last day of Free and first day of FR1 were analyzed. Consistent with our hypothesis, mice took an average of 230 pellets (14.7Kcal) in the Free day and 170 pellets (10.9Kcal) in the FR1 day (significant effect of task: F(1,190) = 120.6, p<0.001, Fig 4B), approximately a 26% reduction in pellets taken. This is consistent with a prior report using FED3 in these same tasks (23). We also observed a significant effect of study location (F(1,190) = 7.5, p<0.001, Fig 4B, B), highlighting the inherent variability among different research labs and mouse colonies, and the importance of performing multi-site studies when possible.

**Figure 4:**
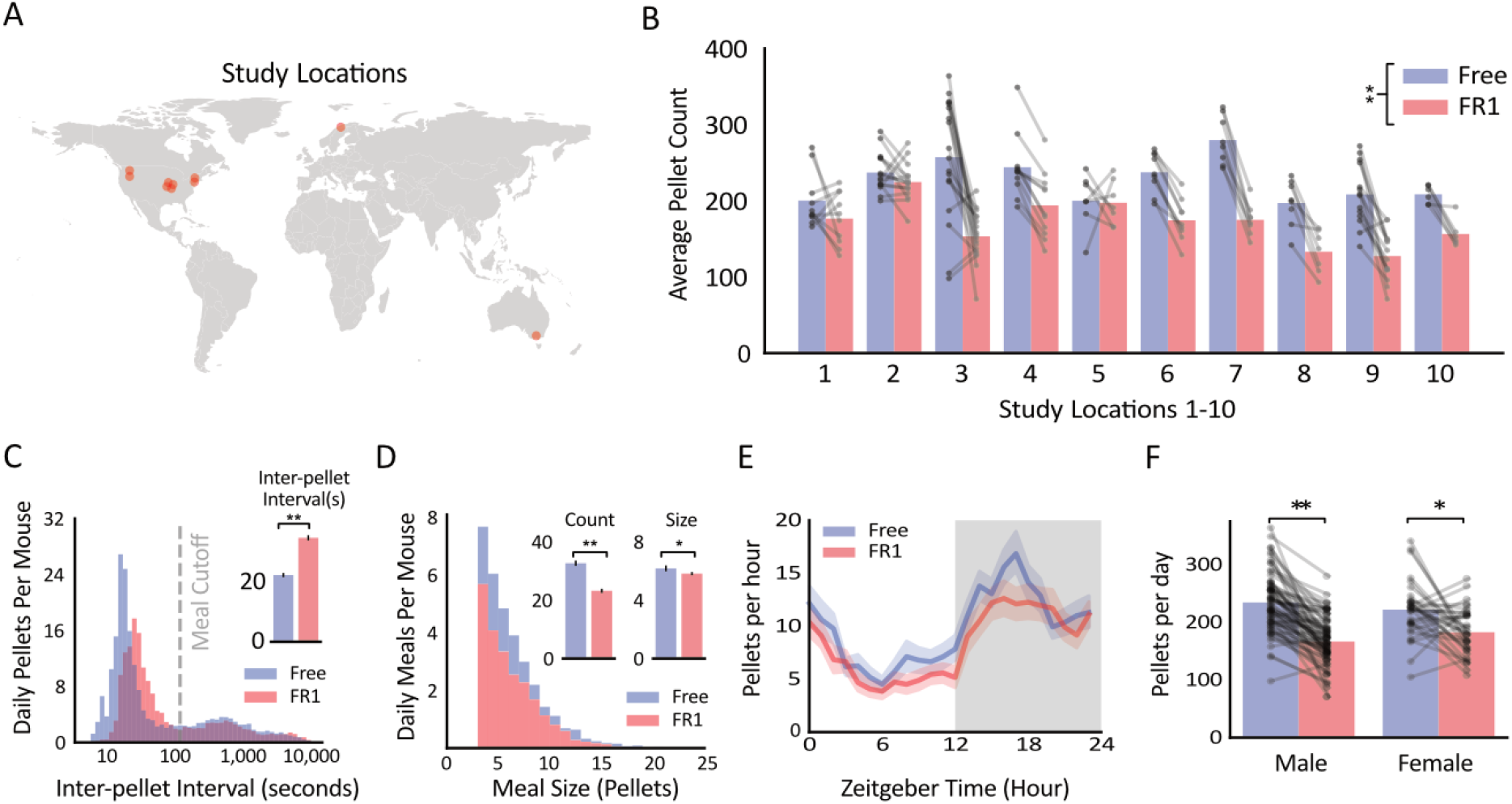
Mice take more grain pellets during Free vs. FR1 access. **A**. Locations of labs from which FED3 data was collected (n=10 study locations). **B**. Average pellet count for mice on the Free vs. FR1 task across study locations (n=105 mice). **C**. Histogram of inter-pellet intervals with a meal cutoff line. (Inset) Average inter-pellet interval for free vs FR1 task. **D**. Histogram of meal sizes for Free vs FR1 tasks (n=105). (Insets) Average count and size of meals in each task. **E**. Circadian plot of average pellets per hour over 24 hours for Free vs FR1 tasks. **F**. Pellets taken by males (n=72) and females (n=33) in Free vs FR1 tasks. ** denotes p<0.001 * denotes p<0.05.

We next examined patterns of pellet taking between the two modes. Meals were defined based on the peaks in the inter-pellet interval distribution (Fig 4C), where pellets within 2 minutes of each other were considered the same meal, and breaks larger than 2 minutes initiated a new meal (22). Mice initiated ∼29% fewer meals on FR1 vs. Free (FR1: 23.2 meals/day, Free: 32.6 meals/day, paired t-test, p < 0.001, Fig 4D), suggesting this effect may account for most of the reduction in pellets taken each day. The average meal size was also ∼0.07% smaller during FR1 than Free (FR1: 5.8 pellets, Free: 6.2 pellets, paired t-test, p = 0.04), and mice took pellets more slowly within meals in FR1 vs Free (FR1: 35.0 seconds between pellets, Free: 22.5 seconds between pellets, paired t-test, p < 0.001, Fig 4C). Differences in pellets taken were not specific to one part of the circadian cycle (Fig 4E) and were observed in both male and female mice (paired t-tests, p<0.005 for both, Fig 4F). Together, we conclude that mice take fewer food pellets when they are required to perform a single nose-poke to obtain them, vs. when the same food pellets are freely available.

## Discussion

We tested the hypothesis that unrestricted and easy access is necessary for high-fat diet induced over-eating, and the corollary that requiring mice to perform small amounts of work to obtain high-fat diet would reduce high-fat diet intake and associated weight gain. We found that introducing a small behavioral requirement – one nose-poke – reduced how much mice interacted with and consumed both high-fat diet and grain-based pellets, and prevented high-fat diet induced weight gain. Critically, the nose-poke is an easy action for mice to complete and there was no limit on how much high-fat diet the mice could earn each day in these experiments. Still, the addition of this small barrier reduced over-eating and almost completely prevented the development of obesity. Our results confirm that in addition to the high level of dietary fat, unrestricted access is necessary for high-fat diet induced over-eating and weight gain in mice. Our results may explain how increasingly easy access to low-cost unhealthy foods has contributed to the obesity epidemic over recent decades (5,7,13), and may inform new solutions for reducing obesity rates by making such foods slightly more difficult to access (24,25).

### Nudging to combat obesity

Strategies that reduce overall food intake are likely necessary to lower obesity rates (2– 4). One idea for how to reduce food intake builds on the “nudge” theory of behavior (26), which posits that small changes in the environment that make specific foods less convenient to obtain can reduce consumption of those foods (27). In 1968, Nisbett demonstrated this effect by offering sandwiches to people of normal weight or people with obesity. The sandwiches were either placed out on a table or in a refrigerator in the room (28). Despite being told they could eat as many sandwiches as they liked, people with obesity ate 37% fewer sandwiches when they were placed in the refrigerator than when those same sandwiches were left out on the table. In this way, making food slightly harder to obtain was effective at reducing intake in people with obesity. A recent meta-analysis determined that changing the position of foods impacted people’s food choices in 16 out of 18 studies analyzed, “nudging” participants towards healthier options by changing their location or proximity to the participant (29). While it is clear that “nudging” can alter food choices and consumption over short durations in laboratory tests, it is less clear if this approach can be used over longer durations to alter body weight (27).

To address this challenge we turned to mice, as we can control their environment and sustain changes in their environment over multiple weeks, which is challenging in humans. We used both an existing pellet-dispensing device (FED3) to dispense grain-based pellets and designed a novel device (the Tumble Feeder) to control access to high-fat diet, to test whether requiring mice to perform an action (one nose-poke) would reduce food intake and weight gain. Surprisingly, requiring mice to perform a single nose-poke greatly attenuated intake of both grain-based pellets and high-fat diets, and reduced high-fat diet induced weight gain, relative to when those same diets were freely available. In this way, our results demonstrate how a small change in the environment can reduce food intake and slow the development of obesity in mice, provided the change is sustained over multiple weeks.

### Out of sight, out of mind: the impact of cue-induced feeding on food intake

A clue to the reduction in intake in our nose-poking task may be gleaned from the temporal structure of their pellet taking. When we broke their daily feeding into “meals”, we found that mice initiated 29% fewer meals when we required them to nose-poke, compared to when the food pellets were freely available. This was the largest change in their feeding patterns and suggests that the direct access to the sight or smell of the food, induced mice to initiate additional bouts of feeding. This may give insight into the neural mechanisms that underlie why mice took less food in the FR1 vs the Free tasks in our experiments.

Several researchers have noted that cues that predict food are sufficient to make animals seek food, even when sated, a phenomenon known as “cue induced feeding” (30,31). Reppucci et al. trained rats to associate a tone with delivery of a food pellet, with control animals receiving the same tones unpaired from food pellet delivery. The researchers then played those tones to the rats in their home-cages for 5 minutes, before giving them the option to eat food pellets or regular laboratory chow for 4 hours. The rats that had learned to associate the tone with the food pellets ate ∼63% more pellets in the consumption test than the control animals, demonstrating that food-paired cues were sufficient to induce over-eating, at least on the time scales recorded in this study. As with auditory cues, the sight or smell of food may serve as additional sensory cues that induce animals to seek and take food in our studies. In this way, keeping food out of sight and not allowing direct contact with the food until the mouse performed a nose-poke may have reduced the salience of the sensory cues from the food and thereby reduced food seeking and taking.

Hunger sensitive hypothalamic arcuate nucleus agouti-related peptide (AGRP) and pro-opiomelanocortin (POMC) neurons are rapidly modulated by the sensory detection of food, which may contribute to subsequent consumption (32). In free-feeding conditions, animals likely investigate the food even when not hungry, simply because they have unrestricted access to it and little else to do in their cages. This may impact the activity of AGRP and POMC neurons to drive further feeding (33,34), in a way that does not happen in the FR1 task. Follow up studies could test the involvement of specific brain circuits or cell types on the behavioral effects we observed. For instance, a brain manipulation that normalized the difference in pellets taken in the Free vs FR1 tasks could inform future brain-based approaches to reduce food intake in food-rich environments.

### Does making high-fat diet harder to get reverse obesity?

We also tested whether requiring a nose-poke to obtain high-fat diet would induce weight loss in obese mice. The obese mice continued to obtain all their daily calories from high-fat diet, but they had to nose-poke once to obtain access to the high-fat diet for one minute each time they wanted to eat. While obese mice did not gain additional weight once the FR1 requirement was introduced, they remained at their elevated body weight and did not lose weight. This highlights the challenges of weight loss, and how obese animals and people will defend their elevated body weights through multiple mechanisms in response to environmental challenges (35,36). Overall, our results suggest that making foods harder to obtain may help prevent, but not reverse, obesity in modern food environments.

## Acknowledgements

This work was supported by R01DK136810 (AVK), R01DK138131 (AVK), the Taylor Family Institute for Innovative Psychiatric Research at Washington University School of Medicine (AVK), the Diabetes Research Center at Washington University (P30DK020579) pilot grants to AVK and MCC, Foundation for Anesthesia Education and Research (FAER) Mentored Research Training Grant (MRTG) GR0033836 (JMT). NIDDK P30 DK56341 to WUSTL Nutrition Obesity Research Center (NORC) and NICHD P50 HD103525 to IDDRC@WUSTL, NIMH R15MH129947 (VAC), Investigator-Initiated Intramural Research Project: 1ZIADK075087-07 (MJK), R01DA049924 (MCC), R01DA058755 (MCC), R01DA056829 (MCC), and Tromsø Research Foundation Starting Grant to JEM (19-SG-JMcC). Indirect calorimetry experiments were completed with assistance from the Diabetes Research Center at Washington University. Thanks to members of the Creed and Kravitz labs, and to Elizabeth Glenn for helpful comments on this manuscript.

## Methods

### Mice and husbandry

C57BL6 mice were individually housed for each experiment on a 12h light/dark cycle. Food was provided as specified in the results and included 20mg grain-based food pellets (Bio-Serv F0163), 20mg sucrose pellets (Bio-Serv F07595), laboratory chow, and 60% high-fat diet (Research Diets #D12492). Water was available *ad libitum* throughout the experiments. All procedures were approved by the Animal Care and Use Committees at Washington University in St Louis, The Artic University of Norway, Monash University, Rutgers University, Temple University, the National Institute of Health, Williams College, Washington State University and Swarthmore College.

### Behavioral testing: Tumble Feeder

The Tumble feeder is an open-source hopper-based device that can be used in the home-cage with mice to control their access to food over multiple weeks.

#### 3D design

The 3D parts for FED3 were designed with TinkerCAD (Autodesk). Tumble Feeders were printed in house with an FDM printer in PLA (Bambu X1 Carbon). We provide editable design files here: https://www.tinkercad.com/things/1IhfIT78Xpz-tumble-feeder-sept2024.

#### Electronics

The Tumble Feeder has two capacitive touch-sensitive nose-poke triggers for the mouse to interact with, a servo-controlled hopper (Analog Feedback Micro Servo with metal gear, Adafruit #1450), touch sensitive bars (UXCELL M2×65mm Stainless Steel Pushrod Connectors) to detect food hopper interactions via capacitive touch sensing, and uses a microcontroller with micro secure-digital (microSD) card slot for controlling tasks, and a Sharp Memory Display (Adafruit #3502), and a DS3231 Precision real-time clock (RTC, Adafruit #3028) for displaying and logging data. The Adalogger M0 contains an ATSAMD21G18 ARM M0 processor that runs at 48 MHz, 256 kB of FLASH memory, 32 kB of RAM memory and up to 20 digital inputs/output pins for controlling other hardware. The microcontroller also contains a battery charging circuit for charging the internal 2200mAhr LiPo battery in FED3, which provides ∼1 month of run-time between charges. The exact battery life depends on the behavioral program and how often the mouse triggers the Tumble Feeder to open as the Servo motor is the main consumer of power. Electronics were programmed in the Arduino IDE and the code is available at https://github.com/KravitzLabDevices/CastleFeeder/.

#### Behavioral tasks

The Tumble Feeder is battery-powered and was placed in the home-cage so mice were able to interact with it around-the-clock for multiple days. The Tumble Feeder was filled with either laboratory chow or high-fat diet, as described in the results, and saved touch times and hopper interaction times on a microSD card for later analysis. The Tumble Feeder was programmed to deliver food based on the task requirements described below:

#### Free-feeding (Free)

In Free-feeding mode in the tumblers were open to allow ad libitum access to food. The touch-sensitive nose-poke triggers had no programmed consequences, and time stamps of touching the metal bars on the hopper were logged to the internal microSD card for later analysis. Every fifteen minutes (96 times per day), the hopper would close and re-open, to control for the noise and movements that occurred in the FR1 session.

#### Fixed-ratio 1 (FR1)

In FR1 mode, the hopper was normally closed and required the mouse to contact the left touch sensor to open it. If the mouse contacted the Left touch sensor the hopper would open for 60s. There were no limits to the number of times the mouse could open the hopper each day, and there were no time-out periods following each time it closed. Time stamps of nose-poke contacts and touching the metal bars on the hopper were logged to the internal microSD card for later analysis.

### Behavioral testing: FED3

The FED3 electronics and hardware have previously been described (Matikainen-Ankney 2021). FED3 is a small battery-powered pellet dispensing device, which was placed in the home-cage so mice were able to interact with it around-the-clock for multiple days. FED3 saves all data on a microSD card for later analysis. There was no limit to the number of pellets the mouse could earn from FED3, which provided 20mg food pellets to mice based on tasks described below. In all tasks, we defined meals as pellets eaten within 2 min of each other, based on inter-pellet interval histograms. In addition, we defined a minimum size of 0.06 g (three pellets) to be counted as a meal.

#### Free-feeding (Free)

In the Free task, FED3 started up with a pellet in its pellet receptacle, and it replaced this with a new pellet 5s after the mouse removed it. FED3 logged the times of all pellet retrieval events for later analysis.

#### Fixed-ratio 1 (FR1)

In FR1 mode, the mouse had to break the left beam-break trigger with its nose (ie: a nose-poke) to cause FED3 to dispense a pellet. After the mouse removed the pellet, it had to nose-poke again to dispense another pellet. There was no time-out period following each pellet delivery, and mice could poke and earn an unlimited number of pellets. However, the nose poke became inactive whenever there was a pellet in the feeding well to prevent buildup of pellets. FED3 logged the times of all nose-poke and pellet retrieval events for later analysis.

### Data analysis and statistics

CSV files generated by Tumble Feeder and FED3 were processed with custom python scripts (Python, version 3.6.7, Python Software Foundation, Wilmington, DE). Data wrangling and plotting used Pandas (37), Matplotlib (38), Pingouin (39), and Seaborn (40) packages. One- or two-way ANOVAs with post-hoc t-tests or paired or unpaired t-tests were used to compare groups as appropriate. p-values < 0.05 were considered significant. Data sets are presented as mean ± SEM. Numbers of animals per experiment is listed as n=number of animals. ChatGPT 4o was used for assistance with coding, but the Authors take full responsibility for all analysis code. All data and analysis and visualization code are available on Github (https://github.com/KravitzLab/Barrett2024).

## References cited

1. USDA ERS - Food Availability (Per Capita) Data System [Internet]. [cited 2024 Aug 19]. Available from: https://www.ers.usda.gov/data-products/food-availability-per-capita-data-system

2. DeSilver D. What’s on your table? How America’s diet has changed over the decades [Internet]. Pew Research Center. 2016 [cited 2024 Aug 19]. Available from: https://www.pewresearch.org/short-reads/2016/12/13/whats-on-your-table-how-americas-diet-has-changed-over-the-decades/

3. Bentley J. U.S. Trends in Food Availability and a Dietary Assessment of Loss-Adjusted Food Availability, 1970-2014. In Unknown; 2017 [cited 2024 Aug 19]. Available from: https://ageconsearch.umn.edu/record/253947

4. Ford ES, Dietz WH. Trends in energy intake among adults in the United States: findings from NHANES. Am J Clin Nutr. 2013 Apr;97(4):848–53.

5. Hall KD. From dearth to excess: the rise of obesity in an ultra-processed food system. Phil Trans R Soc B. 2023 Sep 11;378(1885):20220214.

6. Swinburn B, Sacks G, Ravussin E. Increased food energy supply is more than sufficient to explain the US epidemic of obesity. Am J Clin Nutr. 2009 Dec;90(6):1453–6.

7. Farley TA, Baker ET, Futrell L, Rice JC. The Ubiquity of Energy-Dense Snack Foods: A National Multicity Study. Am J Public Health. 2010 Feb;100(2):306–11.

8. Dunford E, Popkin B. Disparities in Snacking Trends in US Adults over a 35 Year Period from 1977 to 2012. Nutrients. 2017 Jul 27;9(8):809.

9. Qasim A, Turcotte M, de Souza RJ, Samaan MC, Champredon D, Dushoff J, et al. On the origin of obesity: identifying the biological, environmental and cultural drivers of genetic risk among human populations. Obesity Reviews. 2018;19(2):121–49.

10. Elobeid MA, Allison DB. Putative environmental-endocrine disruptors and obesity: a review. Current Opinion in Endocrinology, Diabetes & Obesity. 2008 Oct;15(5):403–8.

11. Baillie-Hamilton PF. Chemical Toxins: A Hypothesis to Explain the Global Obesity Epidemic. The Journal of Alternative and Complementary Medicine. 2002 Apr;8(2):185–92.

12. Cappuccio FP, Taggart FM, Kandala NB, Currie A, Peile E, Stranges S, et al. Meta-Analysis of Short Sleep Duration and Obesity in Children and Adults. Sleep. 2008 May 1;31(5):619– 26.

13. McAllister EJ, Dhurandhar NV, Keith SW, Aronne LJ, Barger J, Baskin M, et al. Ten Putative Contributors to the Obesity Epidemic. Critical Reviews in Food Science and Nutrition. 2009 Dec 10;49(10):868–913.

14. West DB, York B. Dietary fat, genetic predisposition, and obesity: lessons from animal models. Am J Clin Nutr. 1998 Mar;67(3 Suppl):505S–512S.

15. Hariri N, Thibault L. High-fat diet-induced obesity in animal models. Nutr Res Rev. 2010 Dec;23(2):270–99.

16. Deuel HJ, Meserve ER. The effect of fat level of the diet on general nutrition; growth, reproduction and physical capacity of rats receiving diets containing various levels of cottonseed oil or margarine fat ad libitum. J Nutr. 1947 May;33(5):569–82.

17. Tucker LA, Kano MJ. Dietary fat and body fat: a multivariate study of 205 adult females. Am J Clin Nutr. 1992 Oct;56(4):616–22.

18. George V, Tremblay A, Després JP, Leblanc C, Bouchard C. Effect of dietary fat content on total and regional adiposity in men and women. Int J Obes. 1990 Dec;14(12):1085–94.

19. Astrup A, Astrup A, Buemann B, Flint A, Raben A. Low-fat diets and energy balance: how does the evidence stand in 2002? Proc Nutr Soc. 2002 May;61(2):299–309.

20. Hill JO, Melanson EL, Wyatt HT. Dietary fat intake and regulation of energy balance: implications for obesity. J Nutr. 2000 Feb;130(2S Suppl):284S–288S.

21. Yang Y, Smith DL, Keating KD, Allison DB, Nagy TR. Variations in body weight, food intake and body composition after long-term high-fat diet feeding in C57BL/6J mice. Obesity (Silver Spring). 2014 Oct;22(10):2147–55.

22. Matikainen-Ankney BA, Earnest T, Ali M, Casey E, Wang JG, Sutton AK, et al. An open-source device for measuring food intake and operant behavior in rodent home-cages. Cai D, editor. eLife. 2021 Mar 29;10:e66173.

23. Klappenbach CM, Wang Q, Jensen AL, Glodosky NC, Delevich K. Sex and timing of gonadectomy relative to puberty interact to influence weight, body composition, and feeding behaviors in mice. Hormones and Behavior. 2023 May 1;151:105350.

24. Cawley J, Frisvold DE, Hill A, Jones D. The Impact of the Philadelphia Beverage Tax on Purchases and Consumption by Adults and Children [Internet]. Rochester, NY; 2018 [cited 2024 Aug 24]. Available from: https://papers.ssrn.com/abstract=3252029

25. McCarthy M. Soda tax brings sharp fall in sugary drink consumption in Californian city. BMJ. 2016 Nov 4;355:i5940.

26. Thaler RH, Sunstein CR. Nudge: Improving decisions about health, wealth, and happiness. New Haven, CT, US: Yale University Press; 2008. x, 293 p. (Nudge: Improving decisions about health, wealth, and happiness).

27. Rozin P, Scott S, Dingley M, Urbanek JK, Jiang H, Kaltenbach M. Nudge to nobesity I: Minor changes in accessibility decrease food intake. Judgm decis mak. 2011 Jun;6(4):323– 32.

28. Nisbett RE. Determinants of food intake in obesity. Science. 1968 Mar 15;159(3820):1254– 5.

29. Bucher T, Collins C, Rollo ME, McCaffrey TA, De Vlieger N, Van der Bend D, et al. Nudging consumers towards healthier choices: a systematic review of positional influences on food choice. Br J Nutr. 2016 Jun;115(12):2252–63.

30. Reppucci CJ, Petrovich GD. Learned food-cue stimulates persistent feeding in sated rats. Appetite. 2012 Oct 1;59(2):437–47.

31. Weingarten HP. Conditioned cues elicit feeding in sated rats: a role for learning in meal initiation. Science. 1983 Apr 22;220(4595):431–3.

32. Chen Y, Knight ZA. Making sense of the sensory regulation of hunger neurons. BioEssays. 2016;38(4):316–24.

33. Krashes MJ, Koda S, Ye C, Rogan SC, Adams AC, Cusher DS, et al. Rapid, reversible activation of AgRP neurons drives feeding behavior in mice. J Clin Invest. 2011 Apr;121(4):1424–8.

34. Aponte Y, Atasoy D, Sternson SM. AGRP neurons are sufficient to orchestrate feeding behavior rapidly and without training. Nat Neurosci. 2011 Mar;14(3):351–5.

35. Fothergill E, Guo J, Howard L, Kerns JC, Knuth ND, Brychta R, et al. Persistent metabolic adaptation 6 years after “The Biggest Loser” competition. Obesity (Silver Spring). 2016;24(8):1612–9.

36. Guo J, Jou W, Gavrilova O, Hall KD. Persistent diet-induced obesity in male C57BL/6 mice resulting from temporary obesigenic diets. PLoS ONE. 2009;4(4):e5370.

37. Mckinney W. Data Structures for Statistical Computing in Python. Proceedings of the 9th Python in Science Conference. 2010 Jan 1;

38. Matplotlib: A 2D Graphics Environment | IEEE Journals & Magazine | IEEE Xplore [Internet]. [cited 2024 Sep 16]. Available from: https://ieeexplore.ieee.org/document/4160265

39. Vallat R. Pingouin: statistics in Python. Journal of Open Source Software. 2018 Nov 19;3(31):1026.

40. Waskom ML. seaborn: statistical data visualization. Journal of Open Source Software. 2021 Apr 6;6(60):3021.

